# High cholesterol diet modulates macrophage polarization and liver inflammation during early hepatocellular carcinoma progression in zebrafish

**DOI:** 10.1101/299016

**Authors:** Sofia de Oliveira, Ruth A. Houseright, Alyssa L. Graves, Netta Golenberg, Benjamin G. Korte, Veronika Miskolci, Anna Huttenlocher

## Abstract

Diabetes and obesity have been associated with nonalcoholic fatty liver disease (NAFLD)/nonalcoholic steatohepatitis (NASH) and increased incidence of hepatocellular carcinoma (HCC). Here we use optically transparent zebrafish to visualize liver inflammation and disease progression in a NAFLD/NASH-HCC model. We combined a high-cholesterol diet (HCD) with a transgenic zebrafish HCC model induced by hepatocyte-specific activated β-catenin and found that diet induced an increase in liver size and enhanced angiogenesis and neutrophil infiltration in the liver. Although macrophage number was not affected by diet, HCD induced changes in macrophage morphology and polarization with an increase in liver associated TNFα-positive macrophages. Treatment with metformin altered macrophage polarization and reduced liver size in NAFLD/NASH-associated HCC larvae. Moreover, ablation of macrophages limited progression in NAFLD/NASH-associated HCC larvae but not in HCC alone. These findings suggest that HCD alters macrophage polarization and exacerbates the liver inflammatory microenvironment and cancer progression in a zebrafish model of NAFLD/NASH-associated HCC.

## Introduction

Hepatocellular Carcinoma (HCC) is a common cause of cancer-related deaths with increasing mortality worldwide (1). In western societies, 30-40% of HCC patients are obese and have nonalcoholic steatohepatitis (NASH), an aggressive form of nonalcoholic fatty liver disease (NAFLD)(2–5). The development of chronic liver disease in NAFLD has been associated with increased inflammation induced by abnormal lipid accumulation in hepatocytes (6). This aberrant immune activation promotes tissue and organ injury and ultimately leads to fibrosis and carcinogenesis (6). Pro-tumorigenic subsets of neutrophils, macrophages, and other immune cells provide the tumor microenvironment (TME) with growth factors, matrix-remodeling factors and inflammatory mediators that optimize tumor growth (7–10). Hepatic macrophages in particular, including both monocyte-derived or tissue-resident macrophages known as Kupffer cells, have been identified as potential drug targets to treat liver disease (11). Several studies have shown that NAFLD progression to HCC involves inflammatory macrophages(12) and Kupffer cells (13). For example, in a high-fat diet-fed MUP-uPA mouse model, TNF produced by inflammatory macrophages leads to NASH and HCC development in response to hepatocyte ER stress (14). Adaptive immune cells can also be modulators of hepatocarcinogenesis. NAFLD/NASH impairs tumor surveillance by inducing apoptosis of CD4^+^ T cells (15). Taken together, this previous work suggests that the innate and adaptive immune systems are key players in the progression of NAFLD-associated HCC. However, the specific cellular and molecular immune mechanisms that regulate the pathogenesis of early NAFLD/NASH-associated HCC remain unclear.

Several mouse models of HCC are available (16); however, these models have several inherent limitations which make studying temporal and spatial immune interactions with liver cells difficult to characterize *in vivo*(17). To image the immune response in live intact animals, we developed a zebrafish model of NAFLD-associated HCC. Zebrafish have remarkable similarities to humans (80% of genes associated with human disease are present in zebrafish), including hepatic cellular composition, function, signaling, and response to injury (18). Zebrafish combine the complexity of a vertebrate system with unmatched live-imaging capabilites, as the zebrafish liver can be easily imaged in whole intact animals. Moreover, there are already several individual HCC and NAFLD/NASH models established in zebrafish with similar histologic and genetic signatures to human disease (18, 19).

Here we combine a high-cholesterol diet (HCD) with an established transgenic zebrafish model of HCC, which expresses activated *β*-catenin specifically in hepatocytes(20). We find that HCD enhances HCC progression and modulates the immune response in the liver TME. HCD induces changes in macrophage polarization with increased numbers of TNFα-positive macrophages in the liver area. Drug treatment with metformin reduces the numbers of TNFα-positive macrophages and reduces liver size in HCC zebrafish on a HCD. Ablation of macrophages also limits disease progression in HCC larvae on a HCD, but not HCC alone. These findings suggest that HCD alters macrophage polarization and exacerbates the inflammatory microenvironment and progression in a larval zebrafish model of NAFLD/NASH-associated HCC.

## Materials and Methods

### Zebrafish husbandry and maintenance

All protocols using zebrafish in this study were approved by the University of Wisconsin-Madison Institutional Animal Care and Use Committee. Adult zebrafish and embryos up to 5 days post-fertilization (dpf) were maintained as described previously (21). At 5 dpf, larvae were transferred to feeding containers and kept in E3 media without methylene blue. For all experiments, larvae were anesthetized in E3 media without methylene blue containing 0.16 mg/ml Tricaine (MS222/ethyl 3-aminobenzoate; Sigma-Aldrich). Zebrafish lines used are summarized in Supplemental Table 1.

### Diet preparation and feeding of zebrafish larvae

Larval diets were prepared as previously described (22). Briefly, for high-cholesterol diet (HCD), cholesterol (C75209; Sigma) was dissolved in diethyl ether (Sigma) to create a 10% (w/w) solution, then 400 μl was added to 0.5 g of Golden Pearl Diet 5-50nm - Active Spheres (Ingredients: marine fish, krill (23%), fish roe, soy lecithin, yeast autolysate, micro-algae, fish gelatin, squid meal, hydrogenated vegetable fat, vitamins and minerals, antioxidants; Proximate analysis: Protein, 55%; Lipids, 15%; Ash, 12%; Moisture, 8%; Vitamin C, 2550 ppm; Vitamin E, 425 ppm; EPA,10 mg/g; DHA, 12 mg/g). For control normal diet (ND), 400 μl of diethyl ether was added to 0.5 g of GP diet. Diets were left overnight at room temperature to allow ether evaporation. On the following day, diets were ground up into fine particles using a mortar and pestle and stored at −20°C to avoid cholesterol degradation. At 5 dpf, zebrafish larvae were separated into treatment groups in E3 without methylene blue. According to the number of larvae, larvae were maintained in a 15cm petri dish (up to 20 larvae), a small breeding box (20-75 larvae) or a big breeding box (75-up to 150 larvae), and fed for 8 days with ND or HCD (with 2mg, 4.5mg or 6-8mg daily, respectively), E3 was replaced daily. Before any experimental procedure, larvae were fasted for 24 hours. At 13 dpf, larvae were prescreened for HCC (Green Eye marker) or No HCC, and/or liver marker (Green Liver or Red Liver markers) on Zeiss Axio Zoom stereo microscope (EMS3/SyCoP3; Zeiss; Zeiss; PlanNeoFluar Z 1X:0.25 FWD 56mm lens).

### Live imaging

All live imaging was performed using a zWEDGI device as previously described (23). Briefly, an anesthetized larva was loaded into a zWEDGI chamber with the left side down in order to image the left liver lobe. For time-lapse imaging, the loading chamber was filled with 1% low melting point agarose (Sigma) in E3 to retain the larvae in the proper position. Additional Tricaine/E3 was added as needed. All images were acquired with live larvae with the exception of active-caspase 3 staining, Oil Red O staining and macrophage ablation experiments. Images were acquired on a spinning disk confocal microscope (CSU-X; Yokogawa) with a confocal scanhead on a Zeiss Observer Z.1 inverted microscope equipped with a Photometrics Evolve EMCCD camera using a NA 0.5/20× air objective with a 5μm interval. For larvae with large livers, 2×2 tile images were taken. For time-lapse movies of neutrophil and macrophage recruitment, images were taken every 2 min for 40 min. For time-lapse movies of LifeAct activity, images were taken with a NA 0.5/40× air objective, with a 3 μm interval, every 3 min for 2 hours.

### Liver size measurements

To quantify liver size we used three different measurements: liver area, liver surface area and liver volume. To measure liver area, we used double transgenic HCC lines carrying the EGFP liver marker, *Tg(fabp10a:pt*-*β*-*catenin*_*cryaa:venus)/(fabp10a:egfp).* After live imaging, Z series images were reconstructed in 2D maximum intensity projections (MIP) on ZEN pro 2012 software (Zeiss). The area of the liver was measured manually creating a region of interested (ROI) surrounding the larval liver. To measure liver surface area and liver volume, Z series images were 3D reconstructed on Imaris software and a surface of larvarl liver was created manually using the EGFP signal.

### Immune cell recruitment analysis

To quantify leukocyte recruitment we outcrossed double transgenic HCC lines carrying the EGFP liver marker, *Tg(fabp10a:pt*-*β*-*catenin*_*cryaa:venus)/(fabp10a:egfp)*, with a double-transgenic line with labelled macrophages and neutrophils, *Tg(mfap4:tdTomato-CAAX/lyzc:bfp).* For T cell recruitment quantification, we outcrossed HCC line, *Tg(fabp10a:pt*-*β*-*catenin*_*cryaa:venus)*, with transgenic line with labelled T cells, *Tg(CD4.1*-*mCherry/lck:egfp)(24).* After live imaging, Z series images were reconstructed in 2D maximum intensity projections (MIP) on ZEN pro 2012 software (Zeiss). Area of the liver was measured in each larva. Neutrophils, macrophages and T cell were counted within 75 μm from liver. Neutrophil, macrophage and lymphocyte densities were calculated by normalizing the number of immune cells per liver area. Z series images and time-lapse movies were reconstructed on Imaris software using 3D (4D in case of movies) volume rendering modes and 3D/4D reconstructions were used for figures and supplemental movies.

### Vessel density index analysis

To quantify vessel density index we outcrossed the double transgenic HCC line carrying EGFP liver marker, *Tg(fabp10a:pt*-*β*-*cateriin*_*cryaa:venus)/(fabp10a:egfp)*, with a transgenic line with labeled endothelial cells, *Tg(flk1:mCherry)(25).* Images were taken as described above. Z series images were 3D reconstructed on Imaris software and analyzed. Briefly, a surface for the liver area was created using EGFP signal, then a mask for the mCherry signal was generated in order to isolate the signal for vessels in the liver. A second surface was created using mCherry signal in the liver to measure vessel area and volume. The vessel density index was quantified by the following equations:

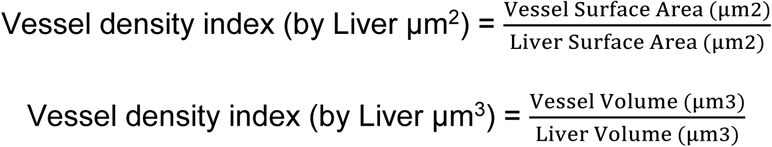

### Cellular and nuclear parameters analysis

In order to evaluate liver disease progression we performed measurements of nuclear:cytoplasmic (N:C) ratio, cell area, cell diameter, nuclear area, nuclear diameter and nuclear circularity. We used a transgenic line with labelled F-actin, *Tg(fabp10a:lifeact*-*egfp)*, outcrossed with transgenic lines with labelled nuclei, *Tg(fabp10a:h2b*-*mCherry)* or *Tg(fabp10a:pt*-*β*-*catenin*_*cryaa:venus)/(fabp10a:h2b*-*mCherry).* All measurements were performed on Z stack images from the first frame of time-lapse movies using the Analysis module of ZEN pro 2012 software (Zeiss). Per larva, 15-30 cells were analyzed and the measurements from those cells were then averaged. Z series images were reconstructed on Imaris software using 3D volume rendering modes and 3D reconstructions which were used for figures and supplemental movies.

### TNFα-positive macrophage quantification

To assess TNFα-positive macrophages in the liver area we outcrossed the HCC line, *Tg(fabp10a:pt*-*β*-*cateriin*_*cryaa:venus)*, with a double transgenic line labelled for macrophages and expressing EGFP under the TNFα promoter, *(Tg(mpeg1:mCherry/TNFα:egfp))(26).* After live imaging, Z series images were 3D reconstructed on Imaris software. Using the Imaris spots tool, total macrophages (mpeg1:mCherry positive cells) were counted within 75μm from liver. TNFα positive macrophages (double positive mpeg1:mcherry/TNFα:egfp cells) were quantified similarly. Images were reconstructed on Imaris software, and 3D reconstructions were used for figures.

### Statistical analysis

All data plotted comprise at least three independent experimental replicates. Least Squared Means analysis in R (www.r-project.org) (27) was performed on pooled replicate experiments, using Tukey method when comparing more than two treatments. This analysis method was used in all experiments with the exception of Lipid and Glycogen accumulation scorings, for which we used Chi-Square test. Graphical representations were done in GraphPad Prism version 6.

## Results

### High cholesterol diet and hepatocyte-specific activated β-catenin increase liver size and induce angiogenesis in both NAFLD and HCC zebrafish larval models

Zebrafish is a powerful model organism for liver disease research, including NAFLD and HCC (18). Here we use a HCC transgenic zebrafish model that expresses hepatocyte-specific activated *β*-catenin (*Tg (fabp10a:pt*-*β*-*cat*))(20). These transgenic zebrafish develop HCC as early as 2 months post-fertilization, and by 6 days post-fertilization (dpf) exhibit features of early HCC such as increased liver size and hepatocyte proliferation(20). We focused our study on the early progression phase of HCC and used 13 days post-fertilization (dpf) *Tg (fabp10a:pt*-*β*-*cat*) larvae, with or without an EGFP liver marker, *Tg(fabp10a:egfp*), referred to here as HCC larvae (Suppl. Table 1). In addition, to induce NAFLD in zebrafish (22, 28), we fed wild type larvae a high cholesterol diet (HCD) from days 5-12 (22), referred to here as HCD larvae. Oil red staining showed that the short term feeding of a HCD was able to induce steatosis in the liver of 13dpf larvae (Fig. S1A). Liver size, a recognized marker for early liver disease progression (20, 29) was increased both with a HCD and in the HCC model (Fig.1A-C; Suppl. Fig.1B and C); importantly these effects were not associated with changes in total body size (Suppl. Fig.1D and E).

**Figure 1:**
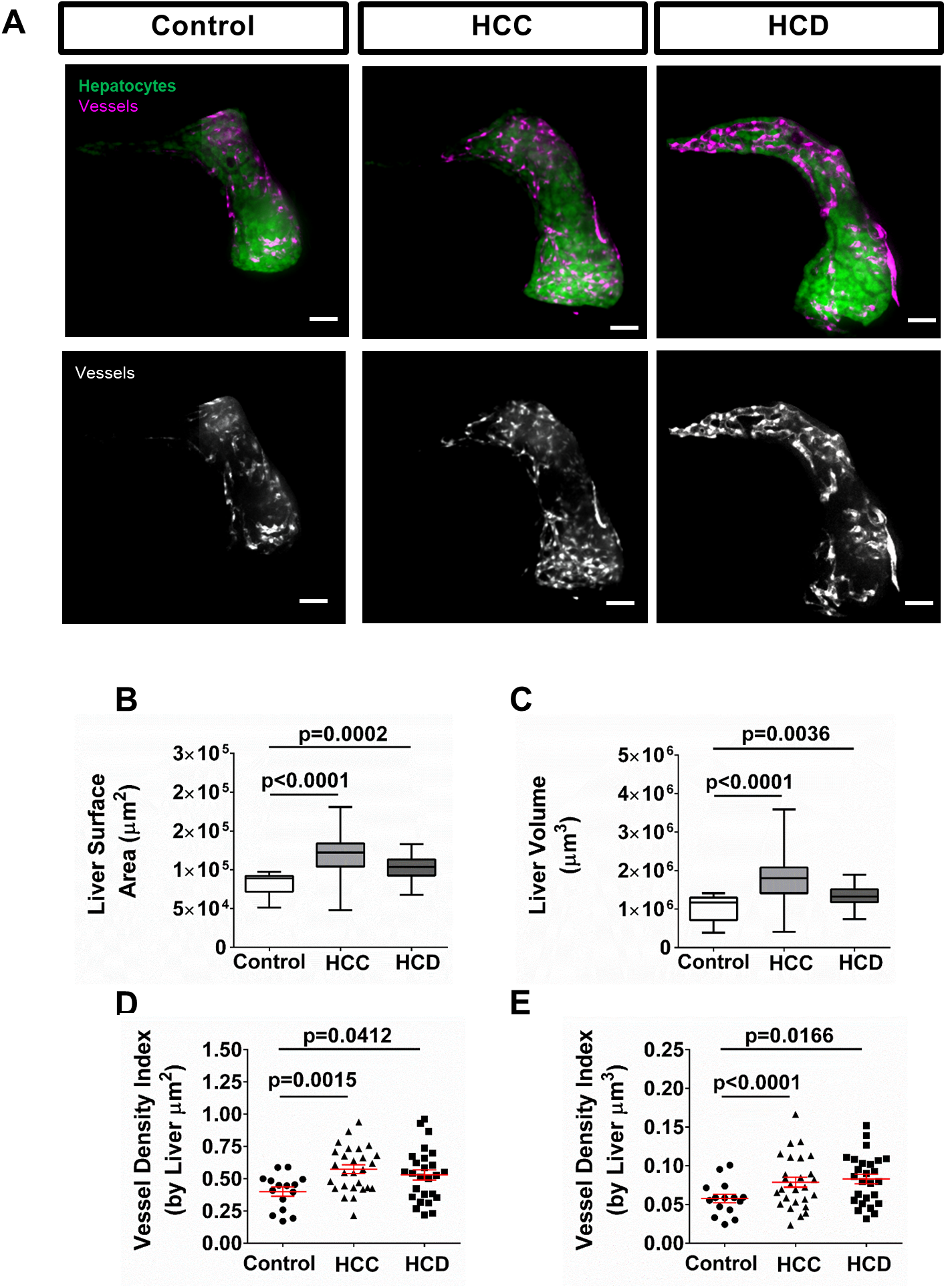
HCD and hepatocyte-specific activation of *β*-catenin promote angiogenesis in zebrafish models. **(A)** Representative 3D reconstructions of livers and vessels in livers of control, HCC and HCD 13-day old larvae. **(B and C)** Box-and-whiskers graphs showing liver surface area **(B)** and liver volume **(C)** in control, HCC and HCD larvae (Control= 15, HCC=27, HCD=26). **(D and E)** Graphs showing vessel density index by liver area **(D)** and vessel density index by liver volume in control, HCC and HCD larvae (Control= 15, HCC=27, HCD=26). All data plotted comprise at least three independent experimental replicates. Dot plots show mean ±SEM, significant p values are shown in each graph.

Increased angiogenesis is an important feature of HCC progression(30), and it has also been associated with NAFLD progression in mouse models after long periods of feeding(31–33). We therefore characterized the vasculature in the liver using an established transgenic line that labels the vasculature, *Tg (flk:mcherry)* (Suppl. Table 1). HCC larvae exhibited increased vessel density index by liver area (Fig.1A and D; Movie S1 and S3) and volume (Fig.1A and E; Movie S1 and S3) as compared to control siblings. Surprisingly, HCD larvae also showed increased vessel density index by liver area (Fig.1A and D; Movie S1 and S2) and volume (Fig.1A and E; Movie S1 and S2) after only a short-term HCD.

### High cholesterol diet and HCC affect liver histologic features differently

Both HCC and NAFLD induce histologic changes in the liver in zebrafish (19, 20, 28). To determine if the characteristic histological changes are present during the larval period, we first performed a blinded, conventional histopathological evaluation of hematoxylin and eosin-stained sections in 13 dpf larvae (Fig. S2A). We performed semiquantitative analysis of lipid accumulation and non-lipid vacuolar change (glycogen accumulation/ballooning degeneration), which are histologic features associated with NASH, and found that these features were more pronounced in HCD larvae (Fig. S2B and C). HCC larvae, but not control or HCD larvae, displayed histologic characteristics of hepatocellular carcinoma, including altered tissue architecture (such as thickened hepatic cords), increased nuclear pleomorphism, and increased mitotic index (Fig. S2D and E). We next took advantage of the optical accessibility of zebrafish larvae to evaluate several cellular and nuclear parameters associated with malignancy using non-invasive imaging techniques. We outcrossed lines with labeled nuclei, *Tg(fabp10a:pt*-*β*-*catenin)/(fabp10a:h2b-mCherry) or tg(fabp10a:h2b*-*mCherry)*, with a line that labels F-actin specifically in hepatocytes, *Tg(fabp10a:lifeact*-*egfp)* (Suppl. Table 1). Using these lines we measured N:C ratio, cell area and diameter, as well as nuclear area, diameter and circularity (Fig. S3). As expected, HCD induced cellular morphology changes characterized by a ballooning effect on hepatocytes, as indicated by increased cell area and cell diameter (Fig. S3). The short-term HCD did not induce nuclear alterations usually associated with carcinogenesis. However, we also observed enlarged hepatocytes with displaced nuclei localized at the edge of the cell in HCD larvae, as reported in NAFLD/NASH disease(34) (Fig. S3). As for the HCC larvae, we observed an increase of hepatocyte area and diameter (Fig. S3). Nuclear alterations associated with carcinogenesis were also observed, such as enlarged nuclei, measured by nuclear area and nuclear diameter, and altered nuclear shape, measured by nuclear circularity (Fig. S3). No significant changes were found in N:C ratio at this stage (Fig. S3). In addition, trinucleated hepatocytes were present in HCC larvae but not in control or HCD larvae (Fig. S3) matching what was found in histologic analysis (Fig. S2F). Our findings suggest that HCD and HCC differently affect histologic features. Nevertheless, our NAFLD and HCC models exhibit histologic features that are characteristic of each human disease.

### NAFLD and HCC zebrafish models exhibit an early increase in leukocyte infiltration

Next, to address the inflammatory response in HCC and HCD larvae, we used a double-labeled macrophage and neutrophil transgenic line (*Tg(mfap4:tdTomato*-*caax)/(lyzc:bfp)* (Suppl. Table 1). The translucency of zebrafish larvae enabled non-invasive time-lapse imaging of leukocytes in the liver area. Both HCC and HCD larvae displayed a significant increase in macrophage influx and density at 13 dpf (Fig. 2A-C; Movie S5-7). Hepatic macrophages were present in control larvae, and displayed crawling and patrolling both in the liver and the surrounding area that was increased in HCC (Movie S5 and S7). Macrophages in HCD larvae displayed a more stationary phenotype than in control larvae (Movies S5 and S6). The HCD also induced a rounder and larger macrophage morphology (Fig. 2A and D; Movie S6), not observed in control or HCC larvae (Fig. 2A and D; Movie S7). Neutrophil influx and density were also increased in HCC and HCD larvae compared to control siblings (Fig. 2A, E and F; Movie S5-7). These results demonstrate that short-term feeding of HCD induces NASH-like phenotype in zebrafish. Altogether, our data suggest that liver inflammation is triggered early both in NAFLD/NASH and in HCC.

**Figure 2:**
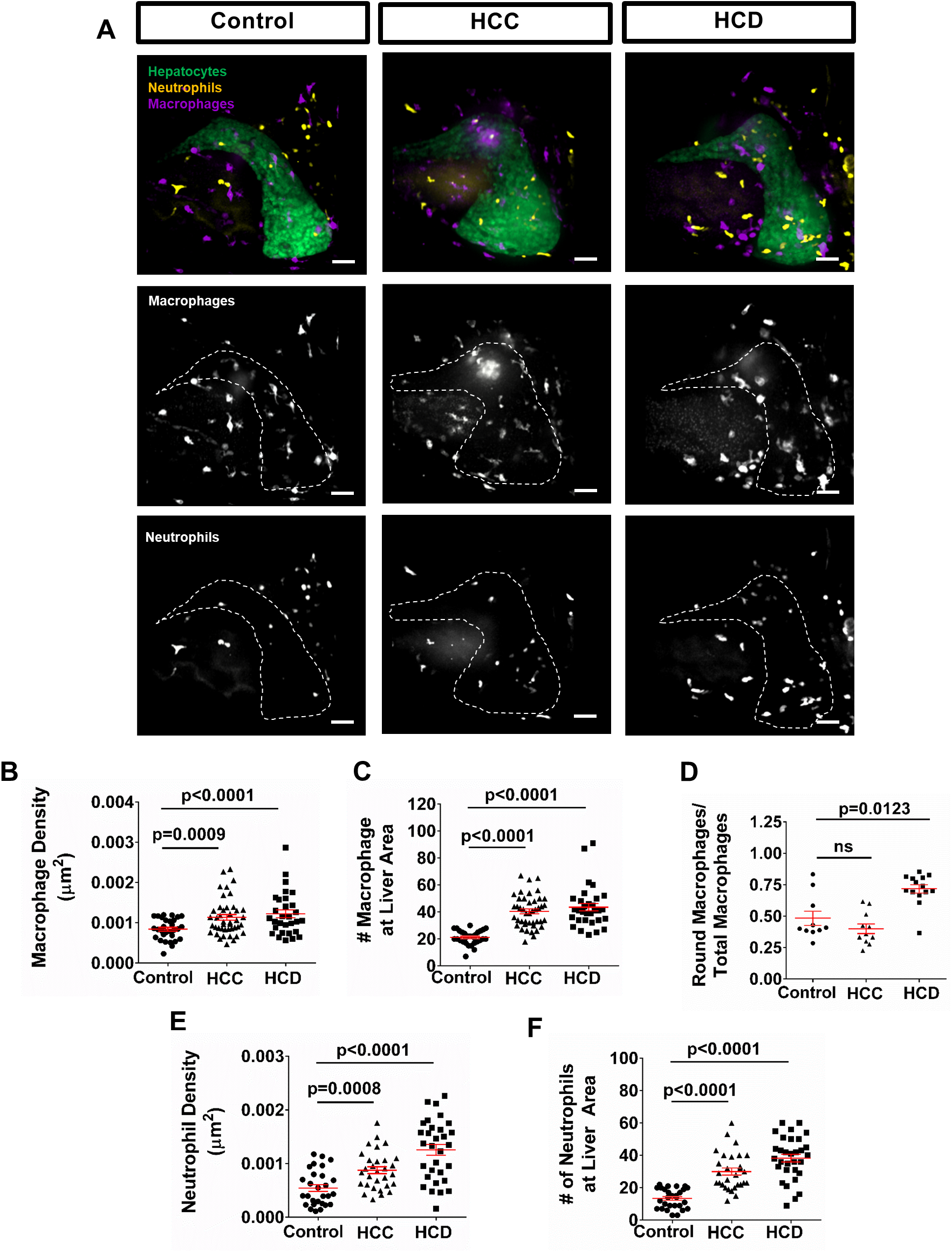
HCD and hepatocyte-specific activated *β*-catenin induce innate immune cell recruitment in zebrafish models. **(A)** Representative 3D reconstructions of liver and leukocyte recruitment to liver area. **(B and C)** Graphs showing macrophage density **(B)** and total number of macrophages **(C)** in liver area in control, HCC and and HCD larvae (Control N=30, HCC N= 43, HCD=30). **(D)** Graph showing ratio of round macrophages over total macrophages at liver area (Control= 10, HCC N= 11, HCD N= 14). **(E and F)** Graphs showing neutrophil density **(D)** and total number of neutrophils **(E)** in liver area in control, HCC and HCD larvae (Control N=28, HCC N= 30, HCD=31). Scale bar= 40μm. All data plotted comprise at least three independent experimental replicates. Dot plots show mean ±SEM, significant p values are shown in each graph.

### High cholesterol diet exarcebates HCC progression in larval zebrafish

NAFLD and NASH can lead to the progression of HCC in humans (1). NAFLD/NASH-associated HCC incidence is increasing, however there is a lack of animal models amenable to live imaging and drug screening. Therefore, we combined our two established models for NAFLD/NASH and HCC, and developed a zebrafish model of NAFLD/NASH-associated HCC by feeding HCC larvae with HCD (HCC+HCD larvae). We first addressed the HCD effect on HCC progression by measuring liver size. In HCC larvae, HCD induced liver enlargement as measured by area, surface area and volume measurements (Fig. 3A-C, S1B and C), without affecting total larval size (Fig. S1D and E). Vessel formation was also increased in HCC+HCD larvae compared to HCC alone (Fig.3A and D; Movie S3 and 4). Importantly, histopathological analysis revealed that HCC larvae fed a HCD displayed features of both NAFLD and HCC (Fig. S4). HCC+HCD larvae also exhibited increased hepatocellular lipid and glycogen accumulation compared to HCC larvae (Fig. S4A, B and C). Although greater nuclear pleomorphism and a trend toward more trinucleated cells was noted in the HCC larvae fed a HCD, significant changes in trinucleated cell numbers and mitotic index were not observed (Fig. S4A, D and E). However, HCD was able to induce cellular and nuclear changes associated with malignancy during early HCC progression phase. HCC+HCD larvae exhibited higher N:C ratio and lower nuclear circularity than HCC larvae (Fig. 3F and G). Live imaging of F-actin dynamics using the Lifeact reporter line (Suppl. Table 1), also revealed an increase in F-Actin activity in hepatocytes in HCC+HCD larvae compared to HCC alone, suggesting a more invasive phenotype (Movies S11-12). Importantly this robus F-actin activity was not observed in control or HCD larvae (Movie S9 and 10). Finally, we also observed an increase in hepatic apoptosis in HCC+HCD compared to HCC larvae using whole-mount immunofluorescence against active-caspase 3 (Act-caspase 3; Fig.3H-J). Together, these findings suggest that HCD is able to enhance malignancy-related histologic and morphologic features in HCC larvae.

**Figure 3:**
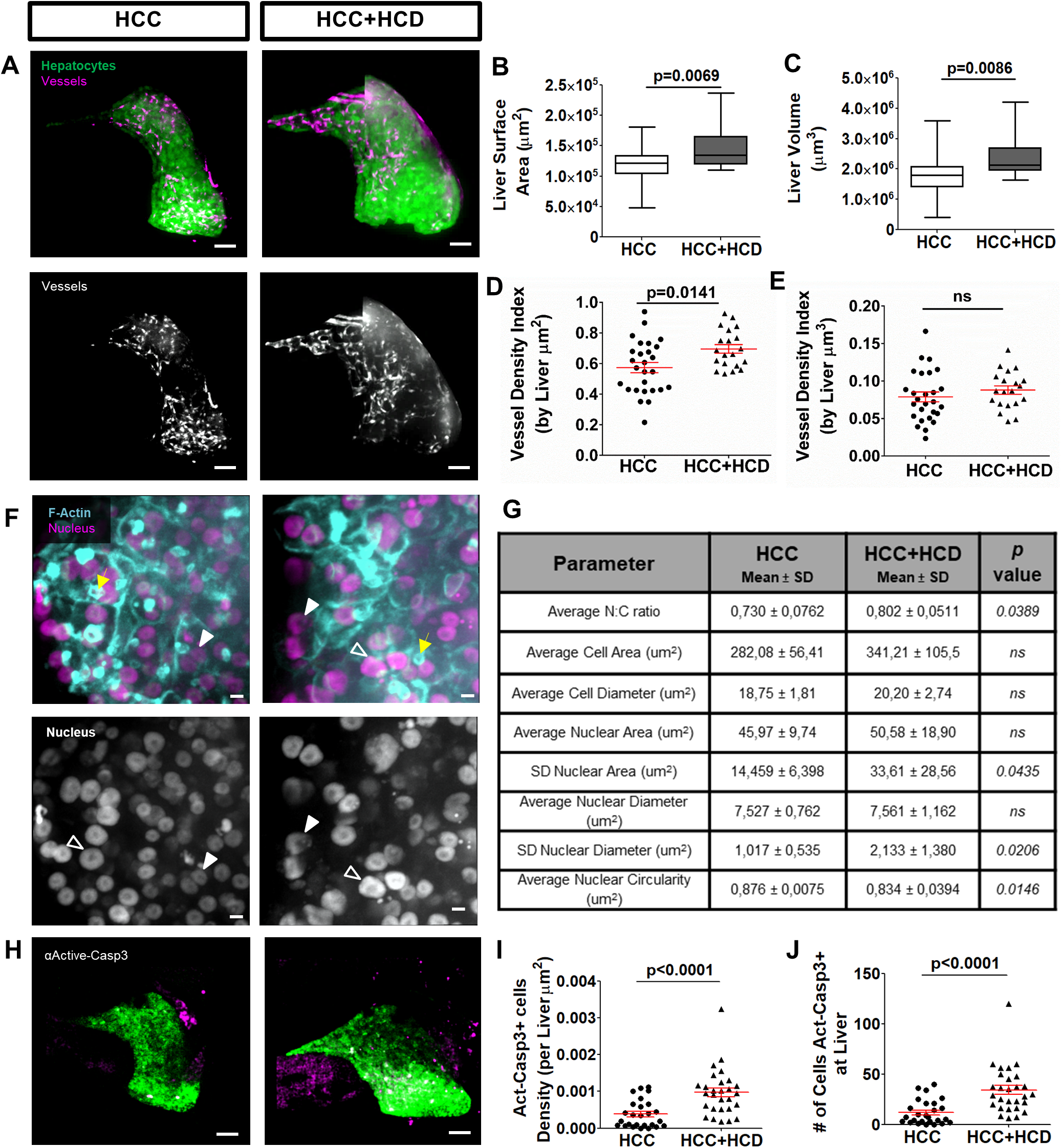
HCD enhances hepatocarcinogenesis in HCC early progression phase. **(A)** Representative 3D reconstructions of livers and vessels in liver of 13-day old HCC and HCC+HCD larvae. **(B and C)** Box-and-whiskers graphs showing liver surface area **(B)** and liver volume **(C)**. **(D and E)** Graphs showing vessel density index by liver area **(D)** and vessel density index by liver volume **(E)** in HCC and HCC+HCD larvae (HCC N=27, HCC+HCD N= 20). **(F)** Representative 3D reconstructions of F-actin and hepatocytes nuclei in liver of in HCC and HCC+HCD larvae. Open arrowheads show Karyomegalic cell; White arrowheads show nucleus with altered shape; and Yellow arrows show dynamic lifeact-positive ring structures. Scale bar 5μm. **(G)** Table with averages of cell and nuclear parameters measured in hepatocytes of HCC and HCC+HCD larvae (15-30 cells/ larvae; HCC N= 8, HCC+HCD N=11). **(H)** Representative 3D reconstructions of liver and Active Caspase 3 in HCC and HCC+HCD larvae. **(I and J)** Graphs showing active-caspase 3 positive cells density in liver **(I)** and total number of active-caspase 3 positive cells in liver **(J)** in HCC and HCC+HCD larvae (HCC N=26, HCC+HCD N= 28). Scale bar= 40μm. All data plotted comprise at least three independent experimental replicates. Dot plots show mean ±SEM, significant p values are shown in each graph.

### NAFLD/NASH-associated HCC larvae display altered immune cell responses

Lipid accumulation in hepatocytes mediates endoplasmic reticulum stress and mitochondrial dysfunction that can also trigger an inflammatory response (6, 35). Therefore, we sought to determine if a HCD alters the immune phenotype of HCC larvae. We found that macrophage influx in the liver was not affected by a HCD in HCC larvae (Fig.4A-C; Movie S7 and 8). However, there was a change in macrophage behavior and morphology, as it was previously observed in HCD larvae (Fig. 2A and D; Movie S6). HCD induced a shift in behavior from the patrolling behavior seen in HCC alone to a more stationary phenotype (Movie S7 and 8). The HCD also induced a rounder and larger macrophage morphology (Fig.5A and C; Suppl. Fig. 3D and E; Movie S8). As for neutrophils, neutrophil influx and density were significantly increased with the combination of HCD and HCC (Fig.4A, E and F; Movie S8). Adaptive immune cells also play an important role in hepatocarcinogenesis (36, 37). To further characterize the immune response in this NAFLD/NASH-associated HCC model, we used a transgenic line with labeled T cells, *Tg(cd4.1:mCherry)/(lck:egfp)* to compare the effects of a conventional diet and HCD (Suppl. Table 1). Surprisingly, in HCC+HCD larvae, T cell influx was reduced in the liver area in comparison to HCC siblings (Fig.4G-I). Overall, these data show that the inflammatory response is triggered early in HCC, and when combined with a NAFLD/NASH model, the HCD significantly alters the innate and adaptive immune response in the tumor microenvironment during the early progression phase of HCC.

**Figure 4:**
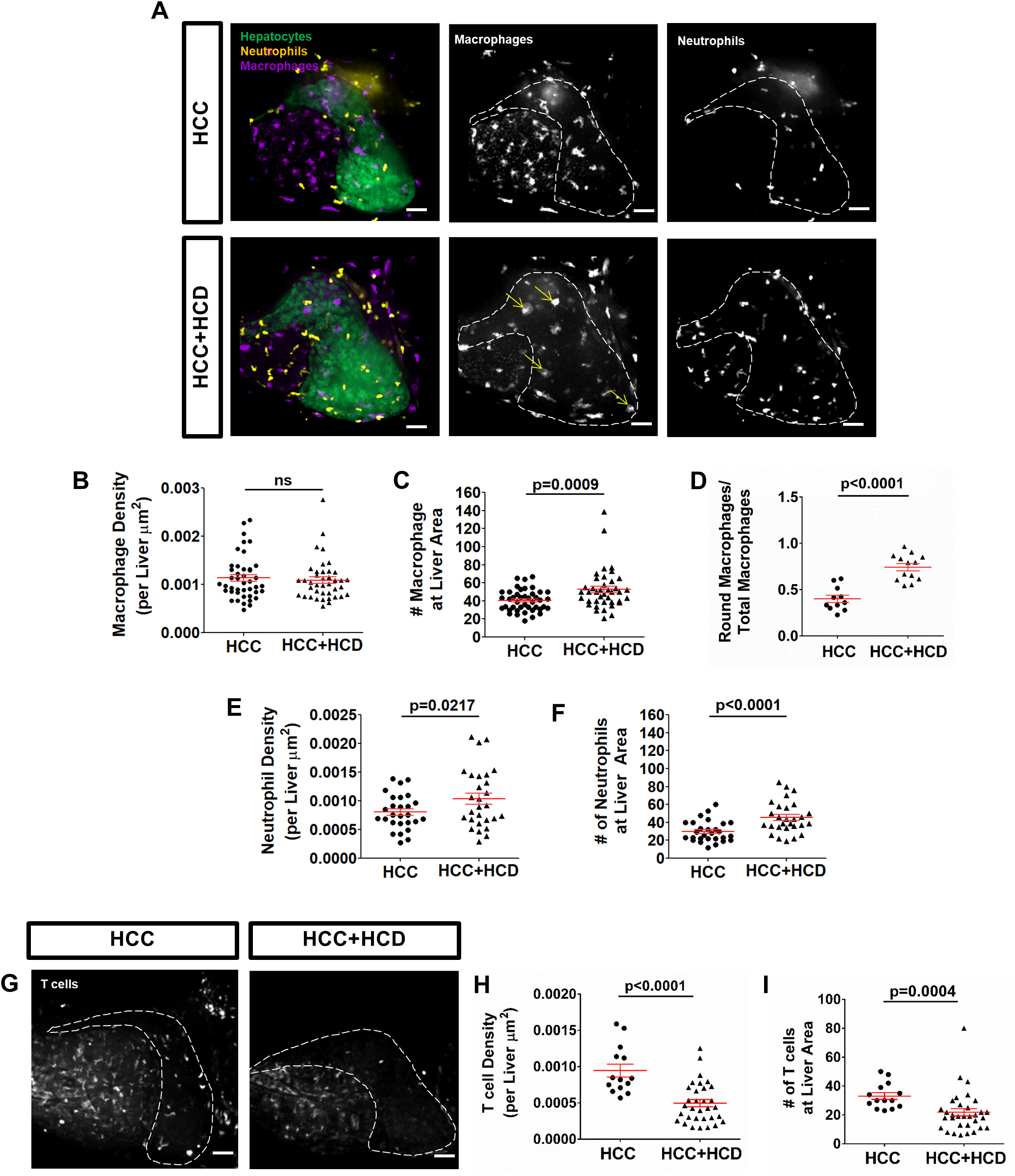
HCD alters the innate and adaptive immune response in zebrafish NAFLD/NASH-associated HCC model. **(A)** Representative 3D reconstructions of livers and leukocyte recruitment to liver area in 13-day old HCC larvae fed with ND (HCC) or HCD (HCC+HCD). **(B and C)** Graphs showing macrophage density **(B)** and total number of macrophages **(C)** in liver area in HCC and HCC+HCD larvae (HCC N=43, HCC+HCD N= 40). **(D)** Graph showing ratio of round macrophages over total macrophages at liver area (HCC N= 11, HCC+HCD N= 13). **(E-F)** Graphs showing neutrophil density **(E)** and total number of neutrophils **(F)** in liver area in HCC and HCC+HCD larvae (HCC N=28, HCC+HCD N= 28). **(G)** Representative 3D reconstructions of lymphocyte recruitment to liver area in HCC and HCC+HCD larvae. **(H and I)** Graphs showing T cell density **(H)** and total number of T cells **(I)** in liver area in HCC and HCC+HCD larvae (HCC N=14, HCC+HCD N= 32). Scale bar= 40μm. All data plotted comprise at least three independent experimental replicates. Dot plots show mean ±SEM, significant p values are shown in each graph.

**Figure 5:**
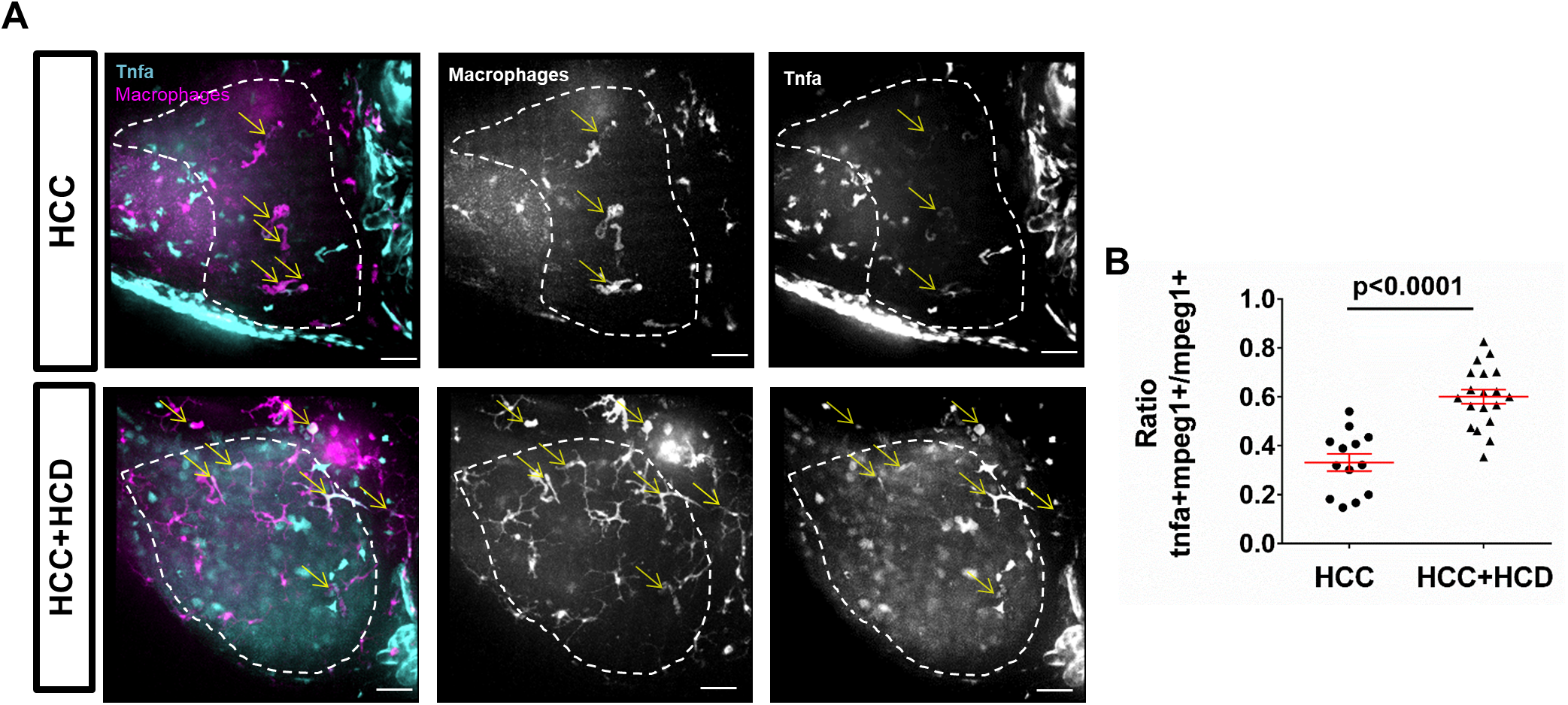
HCD induces macrophage polarization in NAFLD/NASH-associated HCC zebrafish model. **(A)** Representative 3D reconstructions of macrophages and TNFα expressing cells in liver area of 13-day old in HCC and HCC+HCD larvae. Yellow arrows show TNFα-positive macrophages. **(B)** Graph showing ratio of TNFα-positive macrophages over total macrophage number at liver area in HCC and HCC+HCD larvae (HCC N= 13, HCC+HCD N= 19). Scale bar= 40μm. All data plotted comprise at least three independent experimental replicates. Dot plots show mean ±SEM, significant p values are shown in each graph.

### High cholesterol diet induces pro-inflammatory macrophage polarization in zebrafish HCC

Live imaging revealed a change in macrophage morphology and dynamic behavior in HCC larvae in the presence of a high cholesterol diet. Interestingly, we did not observe an increase in macrophage infiltration into the liver with a change in diet. To determine if this change in macrophage morphology is associated with altered polarization of the macrophages we utilized a reporter of TNFα expression, *Tg(tnfa:egfp)* (Suppl. Table 1), to identify pro-inflammatory macrophages in the liver. TNFα is an important marker of macrophage polarization to a pro-inflammatory phenotype, classically referred to as a M1 macrophage, and has been used in zebrafish to identify these macrophage sub-populations (26). It is important to note that other cells in the liver can express TNFα including hepatocytes (38). However, pro-inflammatory subsets of hepatic macrophages are the main source of TNFα in the liver in NAFLD/NASH disease (39, 40). To address macrophage polarization, we crossed the zebrafish HCC model to the TNFα reporter line. We found increased numbers of TNFα-positive macrophages in the HCC+HCD liver compared to HCC control (Fig.5A and B). Taken together, our findings suggest that HCD induces pro-inflammatory macrophage polarization in the liver during early HCC.

### Metformin alters macrophage polarization and HCC progression induced by high cholesterol diet

The finding that HCD induces a change in macrophage polarization in the liver raises the question of whether macrophages are sufficient to induce the changes in liver size and progression induced by a HCD. To address this question we first determined if a drug known to modulate macrophage polarization would alter the phenotype induced by HCD in HCC. Metformin is a drug that induces AMPK activation and is being used to treat diabetes and NAFLD (41–43) and has been shown to reduce HCC incidence and progression in obese/diabetic patients (42). We therefore sought to determine if metformin was able to reduce HCC progression in NAFLD/NASH-associated HCC in larval zebrafish. Interestingly, metformin treatment of HCC+HCD larvae reduced liver size to levels similar to HCC alone (Fig.6A and B), suggesting a reduction of NAFLD/NASH-associated HCC progression. No change was observed in HCC larvae treated with Metformin in the presence of a normal diet (Fig.6A and B), suggesting the effect may be specific to HCD-induced progression. Consistent with prior studies suggesting that metformin alters macrophage polarization *in vitro* (44, 45), we found that metformin treatment reduced the total number and density of TNFα-positive cells in the liver area in HCC+HCD larvae in vivo (Fig.6C and D), suggesting that TNFα polarization of macrophages could play a role in NAFLD/NASH-associated HCC progression.

**Figure 6:**
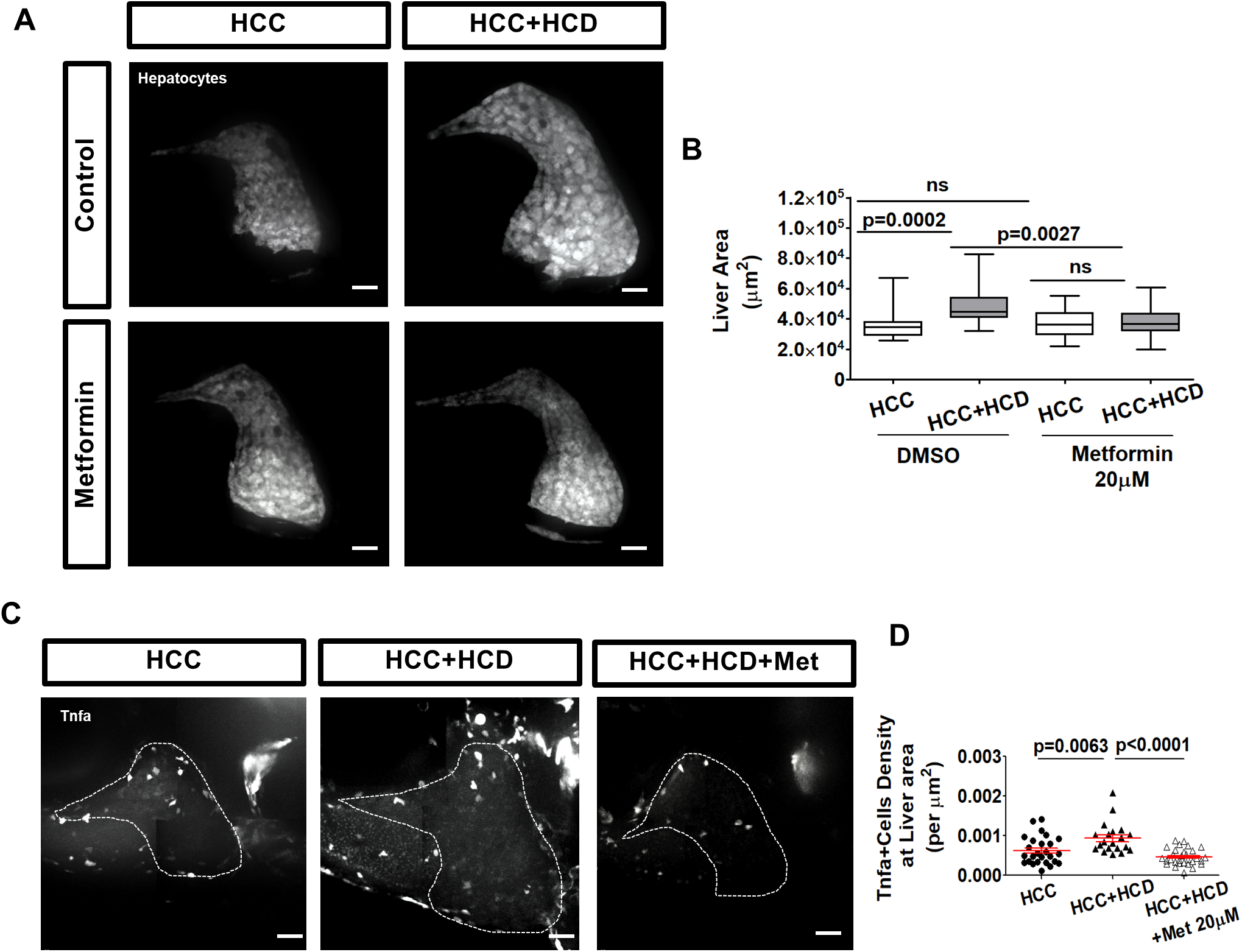
Metformin reduces liver size and TNFα positive cells in the liver in HCC larvae. **(A)** Representative 3D reconstructions of livers in HCC and HCC+HCD larvae with or without metformin treatment. **(B)** Box-and-whiskers graphs showing liver area of HCC and HCC+HCD larvae with or without metformin treatment (HCC N= 22, HCC+HCD N= 27, HCC-Met N= 36, HCC+HCD-Met N= 46). **(C)** Representative 3D reconstructions of TNFα cells in livers from HCC and HCC+HCD larvae treated with or without metformin. **(D)** Graphs showing TNFα positive cell density in livers of HCC and HCC+HCD larvae treated with metformin or control (HCC N= 25, HCC+HCD N= 20, HCC+HCD-Met N= 30). Scale bar= 40μm. All data plotted comprise at least three independent experimental replicates. Dot plots show mean ±SEM, significant p values are shown in each graph.

### Macrophages are necessary for progression of HCC induced by high cholesterol diet

To further address the role of macrophages in the HCC+HCD phenotype, we outcrossed the HCC line with a transgenic line that allows for 80-90% depletion of macrophages, *Tg(mpeg1: NTR*-*eYFP)* (Suppl. Table 1), with metronidazole (MTZ) treatment (46). We began MTZ treatment at 4 dpf to ablate macrophages before we introduced the HCD. Using liver growth as an indicator of disease progression, we found that macrophage ablation reduced liver size to control levels in the HCC+HCD larvae (Fig.7A and B). Surprisingly, macrophage ablation affected liver size in HCC+HCD larvae but not in HCC or HCD alone (Fig.7A and B). Taken together, our findings suggest that macrophages, and specifically macrophage polarization, play a key role in the early progression of NAFLD/NASH-associated HCC in larval zebrafish.

**Figure 7:**
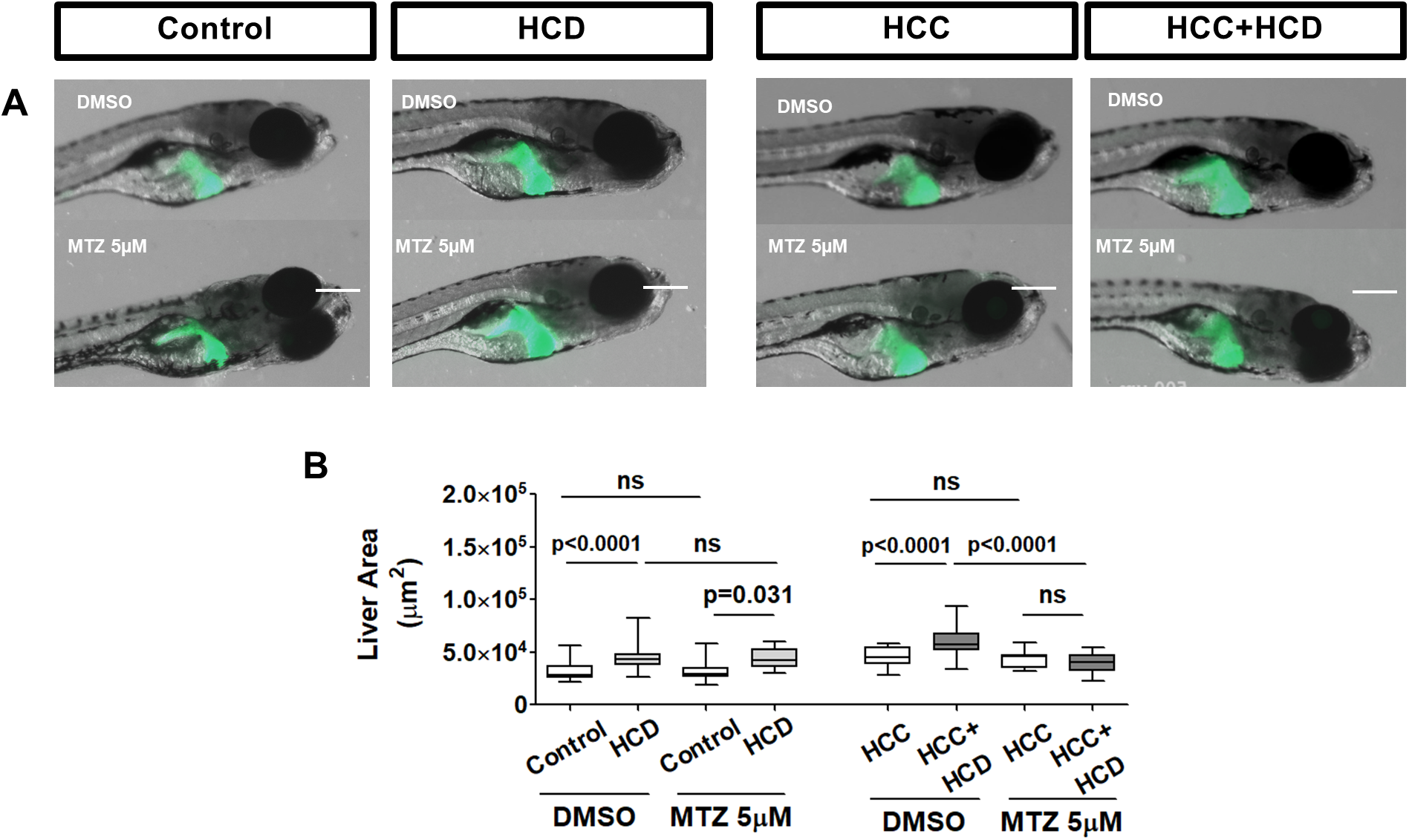
Macrophage ablation affects liver size in NAFLD/NASH-associated HCC larvae but not in HCC or HCD alone. **(A)** Representative images of livers of control, HCD, HCC and HCC+HCD larvae treated with DMSO or metronidazole (MTZ). **(B)** Box-and-whiskers graphs showing liver area in control, HCD, HCC and HCC+HCD larvae treated with DMSO or MTZ (Control-DMSO N= 36, HCD-DMSO N= 44, Control-MTZ N= 20, HCD-MTZ N= 28, HCC-DMSO N= 30, HCC+HCD-DMSO N= 41, HCC-MTZ N= 20, HCC+HCD-MTZ N= 23). Scale bar= 500μm. All data plotted comprise at least three independent experimental replicates. Dot plots show mean ±SEM, significant p values are shown in the graph.

## Discussion

Cancer immunotherapy has brought a new wave of powerful tools to fight cancer. Preclinical data suggests that immunotherapy may play a role in HCC treatment, and currently there are several trials assessing its use in HCC patients (https://clinicaltrials.gov/). Better understanding of immune cell interactions and the molecular players in the liver TME is crucial to identify immunotherapeutics to treat the different types of HCC, including NAFLD/NASH-associated HCC. It is particularly important to understand factors that affect the risk of uncontrolled inflammation that occurs in NAFLD/NASH which can enhance hepatocarcinogenesis. A key gap is the ability to live image intercellular interactions within the liver microenvironment that mediate inflammation and contribute to disease progression. Here we employ tools to visualize features of disease progression and inflammation phenotypes in real time in zebrafish NAFLD/NASH and HCC larval models, and identify a role for macrophage polarization in early progression.

Our findings demonstrate that inflammation and angiogenesis are hallmarks of early liver disease, both in NAFLD/NASH and HCC zebrafish models. A short-term feeding of zebrafish larvae with HCD was enough to induce features of NAFLD/NASH human disease such as liver steatosis associated with hepatocyte ballooning effects, and increased myeloid cell responses characteristic of NASH. Importantly, HCD larvae had increased angiogenesis, consistent with previous reports in other models (31–33). The diet exacerbated the phenotype associated with larval HCC showing a further increase in liver size and angiogenesis. Previous reports have associated increased angiogenesis to NAFLD/NASH (31–33), although as far as we know this is the first time such an effect is being reported in a NAFLD/NASH-associated HCC model and at such an early stage of disease progression. Importantly, HCD also induced alterations in nuclear parameters associated with malignancy during early HCC progression phase. To our knowledge this is the first time that these changes have been observed with such a short duration of treatment, suggesting that short-term dietary changes are enough to enhance hepatocarcinogenesis.

Myeloid cell infiltration has been associated with HCC in humans and in experimental models, but not previously in this zebrafish model of β-catenin induced carcinogenesis or at this stage of hepatocarcinogenesis (20). Both neutrophils and macrophages were recruited to the HCC TME early in development. This pro-inflammatory phenotype is likely an early driver of the malignant progression as supported by our histologic and angiogenesis data. The increase in neutrophil infiltration in the liver both in the presence of HCC and HCC+HCD is consistent with the published literature. Neutrophils can be pro-tumorigenic and enhance hepatocarcinogenesis (47) by releasing growth factors that promote angiogenesis and tumor proliferation (48), or chemotactic cues that modulate recruitment of macrophages and regulatory T cells (49). Increasing neutrophil-to-lymphocyte ratios have been associated with patient’s poor prognosis and are being used as a prognostic factor (50). In addition to the increased liver neutrophil infiltration, NAFLD/NASH-associated HCC larvae also had a decrease in the total number of T cells suggesting that enhanced hepatocarcinogenesis in this model might be associated with decreased tumor surveillance. Interestingly, this effect on T cells is consistent with what was reported by Chi Ma *et al* (15) showing that NAFLD limits liver tumor T-cell surveillance in a mouse model due to reduced numbers of CD4+ T lymphocytes.

Live imaging of F-actin dynamics in larval hepatocytes revealed highly dynamic F-actin structures in HCC larvae but not in control larvae or in the NASH model alone. The dynamic F-actin localized in membrane projections of hepatocytes and was further enhanced in HCC larvae fed a HCD. The increase in F-actin dynamics in hepatocytes may be associated with progression to a more invasive phenotype and an early hallmark of malignant transformation at later stages, and will be informative to follow over time during progression. The relationship between nuclear changes and the dynamics of F-actin structures using long term real time imaging will also likely be informative. Alternatively, these dynamic actin structures may also represent the formation of extracellular vesicles (EVs) such as exosomes (30-150nm), microvesicles (100-1000nm) or apoptotic vesicles (100-5000nm). Hepatocyte-derived EVs have been identified as important mediators for cell-to-cell communications in liver pathogenesis (51). It will be interesting to determine if these changes in F-actin dynamics are involved in the formation of apoptotic vesicles or microvesicles in future investigation.

Our finding that HCD induced changes in macrophage morphology and polarization into a pro-inflammatory TNFα-positive population without affecting overall cell number was particularly intriguing. Indeed, TNFα is a key inflammatory component in NAFLD/NASH and HCC progression (14, 52, 53). This pro-inflammatory cytokine not only serves as a key mediator of hepatocyte apoptosis resulting in liver damage but also plays an important role in cellular proliferation leading to liver regeneration or hepatocarcinogenesis (38). Several cells in the liver can express TNFα, including hepatic macrophages, neutrophils, dendritic cells, natural killer cells, lymphoid cells, endothelial cells and fibroblasts (38, 54). However, pro-inflammatory subsets of hepatic macrophages are the main source of TNFα in liver in NAFLD/NASH disease (39, 40). It has been reported by others that pro-inflammatory sets of macrophages and Kupffer cells play a major role in liver disease progression (55), specially in NAFLD/NASH (12). Macrophage activation in the liver occurs through sensing a wide variety of signals such as HMGB1, ATP, IL1β or ROS that are released from damaged hepatocytes due to lipotoxicity (56). Inflammasome activation in macrophages by danger signals, pathogen-related signals or cholesterol can also amplify the inflammatory response and promote chronic liver inflammation and ultimately HCC (35). In the case of HCC, tumor-associated macrophages (TAM) can play dual functional roles by clearing premalignant senescent hepatocytes and preventing progression to HCC or by providing tumorigenic signals which drive HCC progression (56). Overall, TAM infiltration is generally associated with poor prognosis in HCC and can contribute to HCC progression by promoting proliferation, angiogenesis and later metastasis (7, 11, 56, 57).

In our model we detected a higher number of TNFα-positive macrophages compared to HCC alone, suggesting that an activated pro-inflammatory subset of macrophages exacerbates the inflammatory liver microenvironment and enhances angiogenesis at this early stage of progression. Previous studies reported that depletion of Kupffer cells with clodronate results in the reversal of hepatic steatosis in mice (58). A caveat to our work is that a Kupffer cell marker has not yet been identified in zebrafish (59, 60). However, we and others (59) have visualized a resident macrophage population in the liver that exhibits stellate morphology expected of Kupffer cells. An exciting area of future investigation will be to characterize the role of resident Kupffer cells at this early stage of HCC progression. In our NAFLD/NASH-associated HCC zebrafish model, macrophage depletion also decreased HCC progression. Surprisingly, our data identified macrophages as key modulators of NAFLD/NASH-associated HCC progression in zebrafish but not for HCC alone. It will also be important in the future to determine what macrophage-derived signals are involved in HCC progression to enable potential macrophage-specific drug targets for the treatment of NAFLD/NASH-associated HCC.

In summary, our findings demonstrate that high cholesterol diet alters macrophage polarization and exacerbates the inflammatory microenvironment in the liver, accelerating cancer progression in a zebrafish model of NAFLD/NASH-associated HCC (Fig. 8). Indeed, our findings show that metformin, a drug known to modulate macrophage polarization provides beneficial effects. This supports the power of fish to screen for new small molecules that alter macrophage polarization and may affect progression of HCC. Therefore, this new zebrafish model of NAFLD/NASH-associated HCC is amenable to non-invasive live imaging of the TME and drug perturbations that will further enable the identification of new mechanisms involved in diet induced early HCC progression.

**Figure 8:**
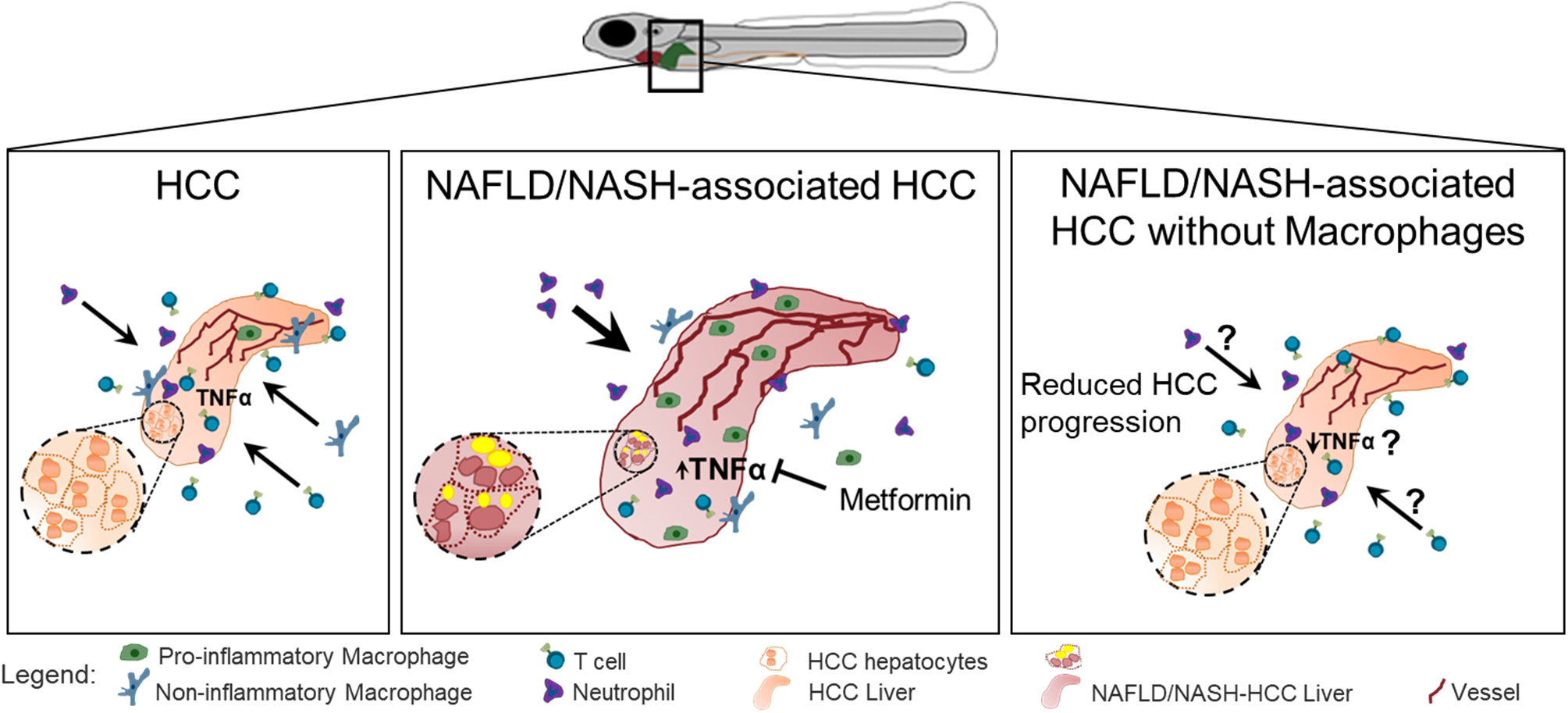
Schematic of HCD early progression in a zebrafish model of NAFLD/NASH-associated hepatocellular carcinoma. Zebrafish HCC larvae show enhanced angiogenesis, exhibit malignancy-associated histologic features and increased immune cell infiltration in the liver TME in early hepatocarcinogenesis. In our NAFLD/NASH-associated HCC zebrafish model, HCD resulted in increased liver size, angiogenesis and neutrophil infiltration. Tumor surveillance is also affected by a HCD characterized by a reduction in T cell density within the liver. Moreover, HCD also induces changes in macrophage polarization characterized by increased numbers of TNFα-positive macrophages. The effect is reversed by Metformin. Importantly, ablation of macrophages reduces disease progression in NAFLD/NASH-associated HCC larvae.

## Acknowledgements

We thank to Dr. Kimberley J. Evason for the zebrafish transgenic *β*-catenin HCC model, to Dr. Kirsten Sadler for the *fabp10a* promoter, to Dr. Randal T. Moon for the *Tg(mpeg*-*NTR*-*eYFP)* line, to Dr. Adam Hurlstone and Dr. David Langenau for *TgBAC(cd4*-*1:mcherry)/Tg(lcK:egfp)* line, Dr. M. Bagnat for the TNFα reporter line *(Tg(tnfα:egfp))*, Dr. Melissa Graham for assistance with telost histopathology, and Dr. Emily E. Rosowski and Dr. Davalyn R. Powell for careful manuscript reading and editing.

Authors Contribution
Conceived and designed experiments: SDO and AH. Performed experiments: SDO, RAH, AG, NG, BGK and VM. Performed analysis: SDO, RAH, AG and VM. Wrote the manuscript: SDO. Critically reviewed and edited the manuscript: RAH, NG, VM and AH

## Abbreviations used in this paper

NAFLD: Nonalcoholic fatty liver disease
NASH: nonalcoholic steatohepatitis
HCC: hepatocellular carcinoma
HCD: high-cholesterol diet
TME: tumor microenvironment
TNF: tumor necrosis factor
dpf: days post-fertilization
ND: normal diet
MTZ: metronidazole
NTR: nitroreductase.

